# Investigating the Functional Role of the DI-DII Linker in Nav1.5 Channel Function

**DOI:** 10.1101/2024.12.01.626264

**Authors:** Emily Wagner, Martina Marras, Shashi Kumar, Jacob Kelley, Kiersten Ruff, Jonathan Silva

## Abstract

The cardiac voltage-gated sodium channel, Nav1.5 initiates the cardiac action potential. Its dysfunction can lead to dangerous arrhythmias, sudden cardiac arrest, and death. The functional Nav1.5 core consists of four homologous repeats (I, II, III, and IV), each formed from a voltage sensing and a pore domain. The channel also contains three cytoplasmic linkers (I-II, II-III, and III-IV). While Nav1.5 structures have been published, the I-II and II-III linkers have remained absent, are predicted to be disordered, and their functional role is not well understood.

We divided the I-II linker into eight regions ranging in size from 32 to 52 residues, chosen based on their distinct properties. Since these regions had unique sequence properties, we hypothesized that they may have distinct effects on channel function. We tested this hypothesis with experiments with individual Nav1.5 constructs with each region deleted. These deletions had small effects on channel gating, though two (430 – 457del and 556 – 607del) reduced peak current. Phylogenetic analysis of the I-II linker revealed five prolines (P627, P628, P637, P640, P648) that were conserved in mammals but absent from the *Xenopus* sequence. We created mutant channels, where these were replaced with their Xenopus counterparts. The only mutation that had a significant effect on channel gating was P627S, which depolarized channel activation (10.13 +/- 2.28 mV). Neither a phosphosilent (P627A) nor a phosphomimetic (P627E) mutation had a significant effect, suggesting that either phosphorylation or another specific serine property is required.

Since deletion of large regions had little effect on channel gating while a point mutation had a conspicuous impact, the I-II linker role may be to facilitate interactions with other proteins. Variants may have a larger impact if they create or disrupt these interactions, which may be key in evaluating pathogenicity of variants.

## Introduction

Intrinsically disordered regions (IDRs) of varying length and function have been identified in the subunits of numerous ion channels including transient receptor potential (TRP)^1^, large conductance Ca^2+^- activated K^+^ channel (BK)^2^, voltage-gated Ca^2+^ (Cav)^3^, ATP-sensitive K^+^ (KATP)^4^, and voltage-gated Na^+^ (Nav)^5^ channels. Due to their lack of a stable tertiary structure and transient protein-protein interactions, disordered regions of ion channels have been largely neglected, and comparatively little is known about their role in channel function.

The cardiac voltage-gated sodium channel, Nav1.5, which is essential for initiating the cardiac action potential (AP), contains multiple segments with predicted disorder.^6^ Activation of Nav1.5 allows Na^+^ ions to rapidly enter the myocyte and depolarize its membrane potential, which leads to myocyte contraction. Nav1.5 dysfunction can lead to dangerous arrhythmias, sudden cardiac arrest, and death. Moreover, variants that pathologically alter Nav1.5 function are associated with inherited arrhythmias such as the long QT syndrome type 3 (LQT3), atrial fibrillation (AF), and Brugada syndrome (BrS). In the case of LQT3, an increase in sodium current prolongs the AP, which can cause proarrhythmic early afterdepolarizations.^7–9^ In contrast to LQT3, BrS1 often involves a reduction in sodium current and slowed conduction as a result, reducing the length of a path required for a re-entrant arrhythmia.^9^ In some cases, these defects can be rescued by antiarrhythmic drugs but individual mutations can vary widely in how responsive they are to this treatment.^10^ Nav1.5 is also found in the gastrointestinal (GI) tract and contributes to the regulation of slow waves and the resting membrane potential and has been implicated in GI pathologies.^11–15^ Variants that alter conductance, expression, and/or mechanosensitivity have been heavily implicated in irritable bowel syndrome (IBS).^16–20^ Some variants have also been implicated in both GI and cardiac disorders.^21^

Nav1.5 comprises four homologous repeats (I, II, III, and IV), encircling a pore region and connected by three cytoplasmic linkers (I-II, II-III, and III-IV).^22,23^ Each repeat includes six transmembrane α-helices (S1 – S6), which are associated with different structural and functional properties. S1, S2, S3, and S4 form the voltage sensing domain (VSD) with the positively charged S4 helix acting as the sensor, moving outwards upon depolarization of the membrane (Fig. 1a). S5, S6, and their large P-loop linker form the pore and Na^+^-selective filter. Each repeat is associated with different functions. Initial pore dilation is caused by conformational changes in I, II, and III. The S4 helices of these domains activate quickly upon membrane depolarization, while that of IV rises after a delay. As the S4 helix of IV rises, an IFM motif in the III-IV linker immobilizes the S4 helices of III and IV in the activated position, facilitating inactivation of the channel and preventing it from conducting sodium ions, even if the membrane is depolarized again.^24,25^ Complementing this tertiary structure are several auxiliary beta subunits^26^ and an alpha-alpha dimerization interaction that is potentially mediated by 14-3-3.^27^ While most of the channel, including the III-IV linker, has been resolved in multiple structures, the I-II and II-III linkers have remained conspicuously absent.^28,29^ In fact, they often must be removed before a structure can be obtained, which suggests that they could be disordered. In further support of this, our preliminary analysis using a well-known algorithm, IUPRED2A, predicted a much higher propensity for disorder in these two linkers than for any other region of the channel (Fig. 1b).

**Figure 1.**
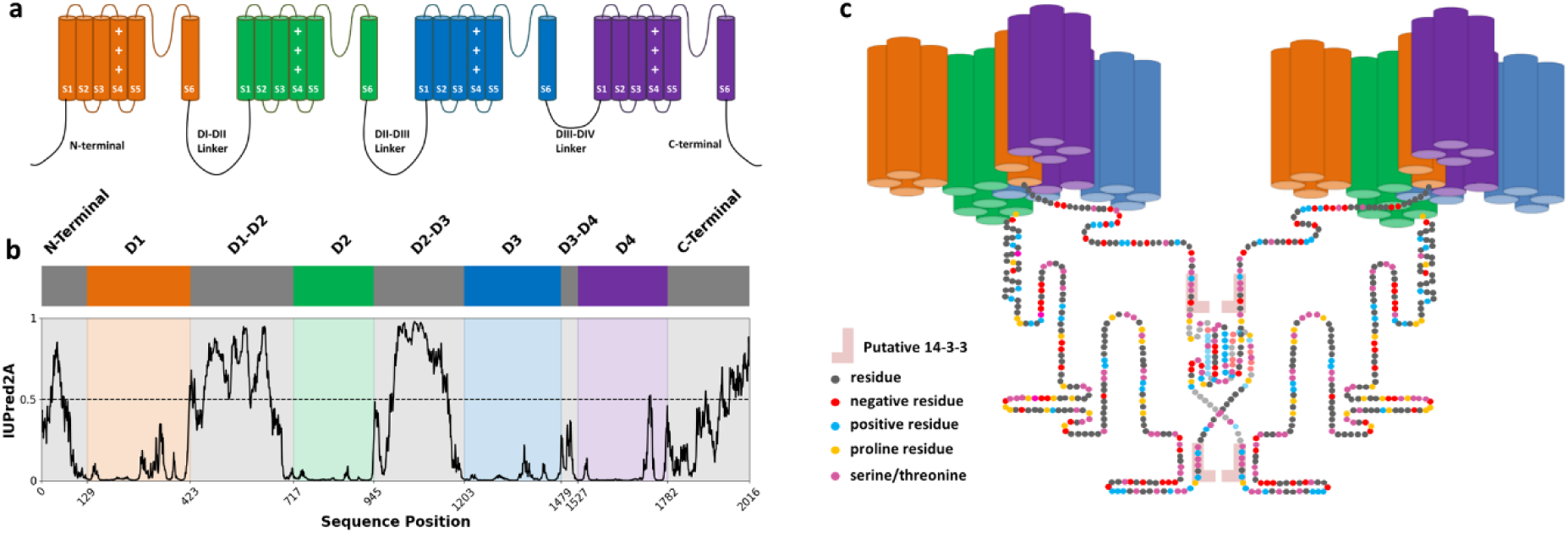
Nav1.5 and the DI-DII Linker. (a) Typical representation of channel topology. (b) Nav1.5 IUPred2A scores, which use an energy estimation method to predict the disorder propensity of each residue in the sequence. (c) DI-DII linker, with putative dimerization and 14-3-3 interactions.

Many variants have been identified in the I-II and II-III linkers, associated predominantly with LQT3 (narrow peaked or biphasic T wave^30,31^), BrS1 (coved-type ST-segment elevation^32^), and AF.^33–36^ Yet their effect on channel function and overall impact on patient health are extraordinarily difficult to predict. Though the linkers have not been extensively investigated, one study created Nav1.4 and Nav1.5 constructs where the I-II and II-III linkers were switched between the isoforms.^37^ The main channel function affected was activation, as channels containing the Nav1.5 linkers exhibited more depolarized activation than channels containing the Nav1.4 linkers. Adding Nav1.5 linkers to Nav1.4 depolarized activation by 6 – 7 mV for single linker substitution and approximately 20 mV for double linker substitution. Adding Nav1.4 linkers to Nav1.5 hyperpolarized activation by 7 mV for single substitutions and 10 mV for double substitutions. Additionally, a previous study^38^ looked at the linkers’ relationship to the effect of papain on the channels. Papain removes fast inactivation from both Nav1.4 and Nav1.5, but only alters the activation of Nav1.5 (∼21 mV hyperpolarizing shift). It was found that each Nav1.5 linker (I-II and II-III) appeared to be responsible for a smaller effect, while both were required for the full shift, when present in either Nav1.4 or Nav1.5. Replacing them in Nav1.5 with those of Nav1.4 eliminated the effect. Notably, though the II-III linkers for Nav1.4 and Nav1.5 are approximately the same size, the I-II linkers are not. The Nav1.4 I-II linker is nearly half the size of the Nav1.5 I-II linker. A portion at the beginning and end are preserved between the two isoforms but large sections from the middle of the Nav1.5 linker are absent in Nav1.4.

Other studies have focused on individual variants and interaction sites. When pathogenic mutations have been identified, they often depend on situational or environmental factors, with many becoming a problem only under certain conditions or in the presence of other molecules or mutations.^39–43^ Though certainly understudied, the I-II linker has been periodically implicated in several critical channel functions. Variants found in the DI-DII linker are associated with disparate effects such as altering mechanosensitivity^18^, disrupting phosphorylation^44^, altering inactivation gating^40,45^, and mitigating the harmful effects of another mutation^46^. The I-II linker has also been linked to various interactions including NEDD4L ubiquitination^47^, ER-retention^48^, 14-3-3 binding^27^, and channel dimerization.^27^ The linker may be heavily involved with post-translation modification, having the largest number of known phosphorylation sites of any region in the channel^49^, including some notable CaMKII^50^ and PKA^44,48,51^ sites, as well as methylation^52^ and ubiquitination sites.^47^ However, the definitive role of the linker is still not well understood and, as a result, attempts to consistently predict the pathogenicity of its variants have so far been largely unsuccessful. In this study, we investigated the basic functional role of the I-II linker and how it regulates the channel by basing our experimental direction on a detailed analysis of its sequence. We theorized that we could tie regions within the linker to specific channel functions, which could provide insight into the mechanisms by which variants promote arrhythmia.

## Methods

### Molecular Biology

cDNAs for human Nav1.5 (Q14524-1 R1027Q) and Navβ1 were produced with the pMAX vector and pBSTA plasmid, respectively. Proline point mutations were made in the Nav1.5 alpha subunit with the QuikChange II site-directed mutagenesis kit (Agilent) and primers from Sigma-Aldrich. Deletions were created from the same Nav1.5 pMax vector through services offered by VectorBuilder. Plasmids were isolated with NucleoSpin plasmid miniprep kit (Macherey-Nagel) and sequenced with Genewiz. They were linearized with the PacI restriction enzyme and purified with the NucleoSpin Gel and PCR Clean-up kit (Macherey-Nagel). Finally, mRNA synthesized from the linearized DNA with the mMessage mMachine T7 Transcription Kit (Life Technologies) and purified via phenol-chloroform extraction.

### Xenopus Oocyte Cut-Open Vaseline-Gap Voltage-Clamp

Before injection, all mRNA was diluted to a concentration of ∼1 ug/uL. Each cell was injected with 50 – 56 ng at a ratio of three Nav1.5 alpha subunits to one Nav B1 subunit and incubated at 18C for 1 – 2 days in ND93 solution (93 mM NaCl, 5 mM KCl, 1.8 mM CaCl_2_, 1 mM MgCl_2_, 5 mM HEPES, 2.5 mM Na-pyruvate, and 1% penicillin-streptomycin, pH 7.4). After incubation, cut-open Vaseline-gap (COVG) voltage-clamp recordings were performed with CA-1B cut-open amplifier (Dagan Corporation). Temperature was maintained at 18 – 19 °C with the HCC-100A temperature controller (Dagan Corporation). External recording solution (25 mM NMDG, 90 mM Na-MES, 20 mM HEPES, and 2 mM Ca-MES2, pH7.4) was used in the guard and recording chambers. Internal recording solution (105 mM N-methyl-D-glucamine (NMDG), 10 mM 2-(N-Morpholino)ethanesulfonic acid (MES) sodium salt (Na-MES), 20 mM HEPES, and 2 mM EGTA, pH7.4) was used in the current injection chamber. This setup was interfaced to a computer with the Digidata 1440A A/D converter (Molecular Devices), where the Clampex software package (v10, Molecular Devices) was used to control experiments and collect data for three voltage protocols. The membrane capacitance was compensated for each recording and P/-8 leak subtraction was performed. The microelectrodes used were pulled from borosilicate capillary tubing to a resistance between 0.2 and 0.5 MΩ.

The activation protocol was used to generate both the conductance-voltage activation curves and the peak current data. For this protocol, currents were recorded during 100-ms interval where the membrane potential was stepped from a holding potential of −120 mV to a test potential, ranging from - 120 to 60 mV. Late current values were obtained from the −20 mV traces.

The inactivation protocol was used to generate the steady-state inactivation (SSI) curves. For this protocol, the cells were pre-pulsed for 200 ms from a holding potential of −100 mV to a conditioning potential, ranging from −150 to 20 mV. Following this conditioning, the membrane potential was then stepped to −20 mV for 20 ms to record channel availability after inactivation. Both the activation and SSI curves were plotted as function of membrane potential and fit with a Boltzmann equation:

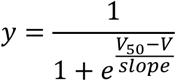

The recovery protocol was used to generate the recovery from inactivation data. For this protocol, a pre-pulse of 200 ms (P1) and a test pulse of 20 ms (P2) of −20 mV were applied. These two pulses were separated by a period where the membrane potential was held at the holding potential of - 120 mV for varying durations, ranging from 1 to 1000 ms. The recovery from inactivation was plotted as a function of time and fit with a double exponential equation:

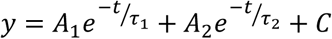

### Electrophysiological data analysis

Electrophysiological data was analyzed with a custom Python package, which was initially designed to recreate and streamline the basic functionality of the Clampfit analysis package (Molecular Devices) but was further optimized for sodium channel data analysis. Variant/deletion channel results were only compared to wild-type results collected on the same day from the same batch of oocytes. Statistical significance was evaluated using the two-tailed t-test.

### Sequence Analysis

IUPred2A scores were obtained from IUPred2A web server that uses the method described by Mészáros, Erdös, and Dosztányi.^53^ The Net Charge Per Residue (NCPR) was calculated for all sliding windows of length 5. Then, the mean NCPR per residue was calculated by taking the mean of all sliding windows that contained that residue. Kyle & Doolittle (K&D) Hydropathy was determined by calculating the mean hydrophobicity score within a five-residue window around an individual residue. The hydrophobicity scores for each residue were taken from the published scale.^54^

### Phylogenetic Analysis

A set of sequences was obtained by searching the NCBI Protein database for eukaryotic proteins between the lengths of 1800 and 3000 residues matching the terms “sodium channel protein type 5 subunit alpha”. This set contained representatives from 350 different species. Alignment was performed with Clustal Omega in a multi-step process. First, the full channel sequences were aligned and the portions corresponding to the human DI-DII linker were extracted. These DI-DII linker sequences were then further aligned and portions corresponding to our designated regions in the human sequence were extracted. Finally, these region sequences were aligned, and their proline fractions were calculated.

## Results

### The DI-DII Linker can be split into eight regions with distinct properties

We separated I-II linker into eight regions ranging in size from 32 to 52 residues (**Fig. 2**), chosen based on their distinct properties. The N-terminal (411 – 457) region is directly connected to the structured I domain and the C-terminal (685 – 717) is connected to the II domain. These regions, especially the C-terminal, have low predicted propensity for disorder, as represented by the IUPred2A score. IUPred2A uses an energy estimation method to determine how favorable a residue is towards forming contacts with other residues. Generally, inter-residue interactions stabilize protein structures, so contact-favoring residues are more likely to promote order within a protein and contact-hindering residues, more likely to promote disorder. A low IUPred2A score suggests that a region is likely at least partially, if not fully, ordered.

**Figure 2.**
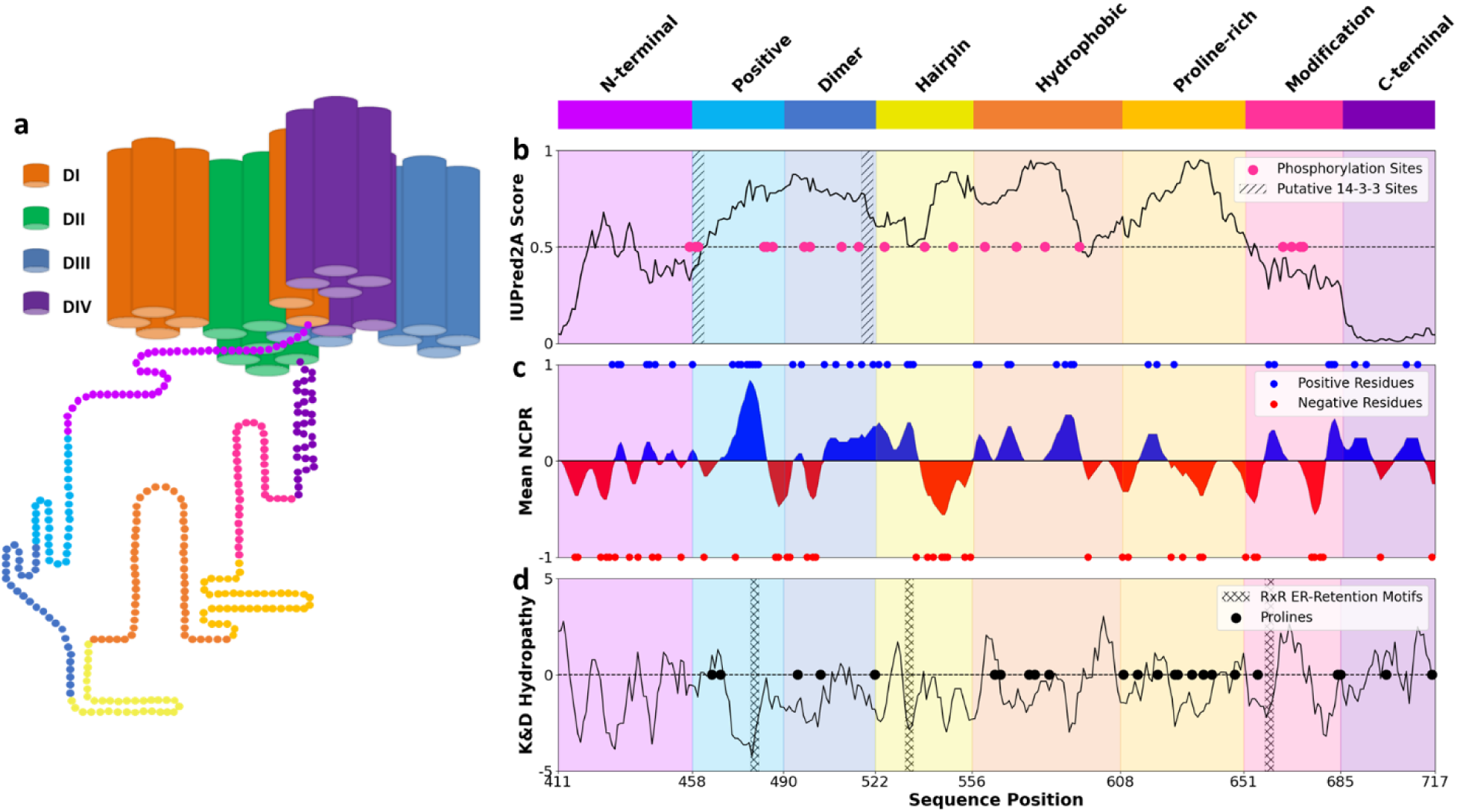
Summary of the properties used to divide the I-II linker into regions. (a) Representation I-II linker attached to Nav1.5 with linker residues color-coded by region. (b) Plots the IUPred2A scores, which use an energy estimation method to predict the disorder propensity of each residue in the sequence. Phosphorylation sites^49^ are notated with pink dots and putative14-3-3 interaction sites^27^, with grey boxes. (c) Mean Net Charge Per Residue (NCPR), calculated for a five-residue window around an individual residue. Positively charged residues are notated with blue dots and negatively charged residues, with red dots. (d) Kyte & Doolittle Hydropathy scores, calculated by taking the mean hydrophobicity score within a five-residue window around an individual residue. Proline residues are notated with black dots.

The regions spanning from residue 458 to 521 form a section implicated in the potential dimerization of the channel^27,55^, consisting of one alpha-alpha and two 14-3-3 interaction sites. Channel dimerization may play a role in targeting and clustering channels at specific locations within the cell.^56^ Additionally, gating is coupled between the two subunits and this coupling appears to be mediated by 14-3-3, suggesting that this region of the linker could play a role in gating. However, these roles and behaviors are currently controversial.^57,58^ The Positive (458 – 489) region starts at the first putative 14-3- 3 interaction site and includes the first half of the section. It contains a large cluster of positively charged residues, including an RxR ER-Retention motif. The Dimer (490 – 521) region contains the second half and ends at the second putative 14-3-3 interaction site. It also contains a notable CaMKII-interaction site, S516.

Following these sections is a region we termed the Hairpin (522 – 555). It exhibits a high degree of charge segregation, with positively charged residues relegated to its first half and negatively charged residues to the second. The distribution of electrostatic charge is often a key determinant of IDP conformational behavior. Typically, disordered proteins with segregated charge patterning adopt hairpin-like conformations due to the attraction between the oppositely charged sides. This region also contains two PKA phosphorylation sites (S525 and S528) that are associated with a peak current increase through the mediation of nearby ER-retention motifs.^44,48^

The hydrophobic (556 – 607) region has two notable spikes in hydrophobicity and a relatively low fraction of charged residues. Disordered proteins tend to be deficient in hydrophobic residues, which promote conformational collapse and the formation of stabilizing, hydrophobic cores. It contains two notable CaMKII-interaction sites, S571 and T594. Previous reports show that S571 may regulate late Na^+^ current. In Glynn et al. 2015, it was found that in mouse ventricular cardiomyocytes, S571E increased late current while S571A decreased it.^50^

The Proline-rich (608 – 650) region has the highest proline content in the linker. In general, proline content is higher in the intrinsically disordered proteome than in the ordered proteome. Prolines tend to promote conformational expansion through their unique relationship with secondary structure. Unlike other amino acids, prolines have a second connection between their side chain and backbone through the backbone nitrogen.^59,60^ This removes a proton, which is present on the amide backbone nitrogen in other amino acids and plays an important role in the formation of most standard secondary structures.^59–61^ As a result, prolines can disrupt secondary structures, promoting disorder. When, however, there are several close in sequence, prolines can form their own helical structure, known as polyproline II (PPII).^62^ PPII may promote disorder from a tertiary structural perspective by locally expanding the backbone, preventing compaction. The effect of prolines on protein conformation can be variable, however. Depending on sequence patterning, for example, prolines have been known to promote compaction in some cases. Prolines also are more likely than other amino acids to undergo cis-trans isomerization^59,60^, which has led in recent years to their study as a potential switch mechanism.^63,64^ Their unusual properties and general unpredictability make them an interesting consideration in the study of IDPs.

Finally, the Modification (651 – 684) region contains an RxR ER retention motif and a notable cluster of phosphorylation sites, investigated in Lorenzini et al. 2021.^49^ In HEK-293 cells, channels with either S664 or S667 mutated to either the neutral residue, alanine, or the phosphomimetic residue, glutamate, showed a marked depolarization in the voltage dependence of their inactivation. Channels with S671 mutated to the phosphomimetic residue, glutamate, showed a decrease in peak current density. Lorenzini et al. suggest that the alteration of channel properties occurs through a disruption of native phosphorylation.^49^ The presence of an ER-retention motif may suggest a further role for this region in controlling channel expression.

Since these regions had distinct sequence properties, we hypothesized that they may have distinct effects on channel function, which was tested experimentally.

### Deleting whole regions had little to no effect on the function of channels expressed in oocytes

We created individual Nav1.5 constructs where we deleted each of the regions, expressed them in *Xenopus* oocytes, and recorded the current with a COVG voltage-clamp for the activation and steady-state inactivation protocols (Fig. 3). We were able to acquire data for the deletion of each of the full regions between the partially structured termini. Interestingly, deletions of the regions identified in the previous section had very little effect on channel function (Fig. 4a-f and Table 1). The Proline-rich region deletion does show a statistically significant (p < 0.05) activation shift (**Fig. 4g**); the Hydrophobic region deletion shows a significant (p < 0.05) SSI shift (**Fig. 4h**); and the Positive region deletion, a significant (p < 0.05) change in the slow time constant of the recovery from inactivation (**Fig. S1a**). However, the small size of these changes (Proline-rich deletion activation: 5.14 +/- 1.70 mV, Hydrophobic SSI: 3.71 +/-1.11 mV, Positive slow tau: 16.16 +/- 4.07 ms) makes their functional relevance uncertain. Noticeably, the effects on the recovery time constant did not seem to have much impact on the recovery plot (**Fig. S1d**). Comparatively, the Hydrophobic region deletion has a statistically significant change in peak current magnitude (**Fig. 4i**) that is quite large (−1.77 +/- 0.69 uA), making it the only deletion that has a high likelihood for relevant functional effects. No statistically significant effects on the fast time constant of the recovery from inactivation (**Fig. S1b**) or the late current (**Fig. S1c**) were seen for any deletion.

**Figure 3.**
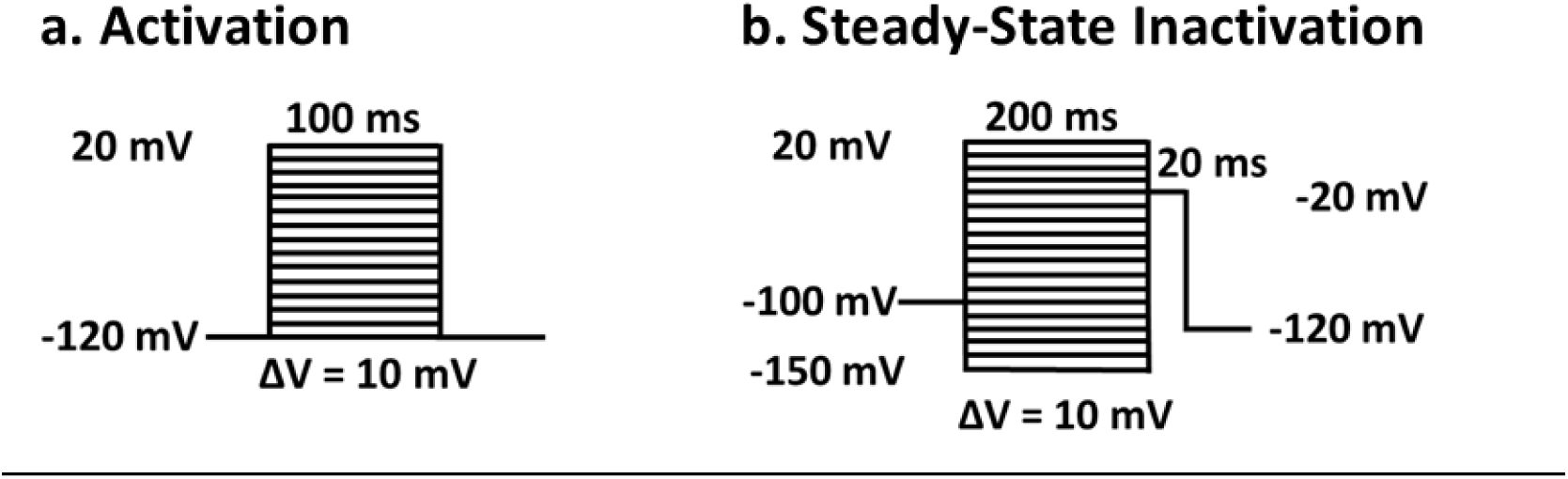
Stimulation protocols used to obtain the (a) activation and (b) steady-state inactivation curves.

**Figure 4.**
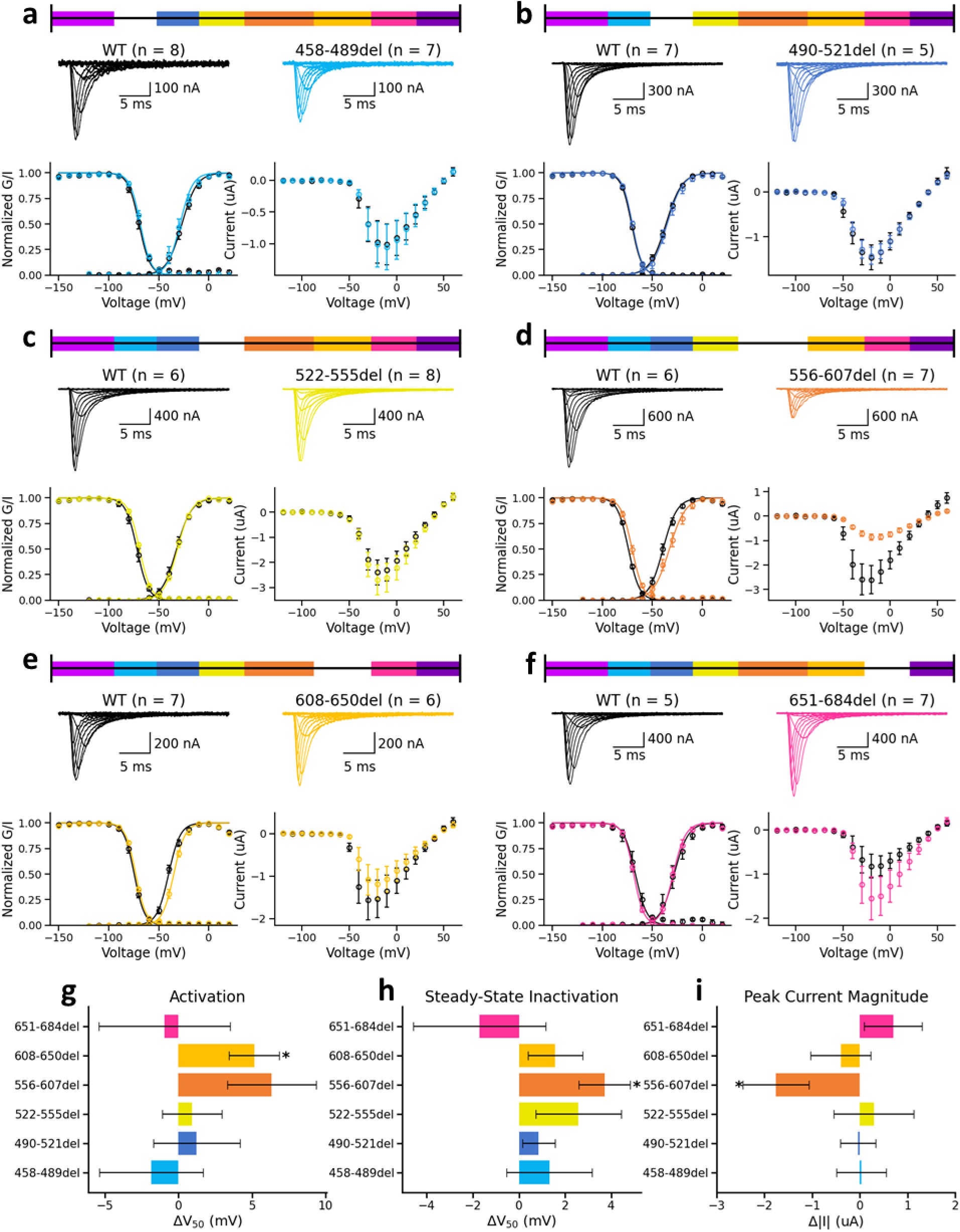
Deleting the different regions has little effect on channel function. Xenopus oocyte data for deletions of the (a)Positive (458 – 489), (b) Dimer (490 – 521), (c) Hairpin (522 – 555), (d) Hydrophobic (556 – 607), (e) Proline-rich (608 – 650), and (f) Modification (651 – 684) regions compared to wild-type data collected on the same days. For each deletion, there are representative traces for the wild-type (black) and deletion (color) channels at the top of the section. Below the wild-type trace, the mean (+/- SEM) normalized conductance (G) and current (I) are plotted as functions of membrane potential and fit with single Boltzmann equations, representing channel activation and steady-state inactivation (SSI), respectively. Below the deletion trace, the mean (+/- SEM) peak current is plotted as a function of membrane potential. This may roughly corresponds to the number of channels that reach the membrane and can be used as an indicator of channel expression. (g) Difference between the means (+/- SEM) of the deletion and wild-type activation V_50_ from the Boltzmann fit (* indicates p-value < 0.05). The Proline-rich region deletion shows the only statistically (p-value < 0.05) significant shift. However, the small size (5.14 +/- 1.70 mV) of this shift makes its functional relevance uncertain. (h) Difference between the means (+/- SEM) of the deletion and wild-type SSI V_50_ from the Boltzmann fit (* indicates p-value < 0.05). The Hydrophobic region deletion shows the only statistically (p-value < 0.05) significant shift. However, the small size (Hydrophobic: 3.71 +/-1.11 mV) of this shift makes its functional relevance uncertain. (i) Difference between the means (+/- SEM) of the deletion and wild-type peak current magnitude (* indicates p-value < 0.05). The Hydrophobic region deletion shows the only statistically (p-value < 0.05) significant shift. This shift is quite large (−1.77 +/- 0.69 uA), suggesting some potential for functional relevance.

**Table 1.** Activation (GV) V_50_, Steady-State Inactivation (SSI) V_50_, and peak current magnitude (Peak, |I|) values for the I-II region deletions.

| Construct | WT GV $V_{50}$ | Variant GV $V_{50}$ | GV $\Delta V_{50}$ (mV) | WT SSI $V_{50}$ | Variant SSI $V_{50}$ | SSI $\Delta V_{50}$ (mV) | WT Peak | Variant Peak | $\Delta \vert I\vert $ (uA) |
| --- | --- | --- | --- | --- | --- | --- | --- | --- | --- |
| 458-489del | -26.88 +/- 1.94 | -28.71 +/- 2.63 | -1.84 +/- 3.52<br>(p = 0.6038) | -69.75 +/- 1.49 | -68.43 +/- 0.88 | 1.32 +/- 1.85<br>(p = 0.5037) | -1.04 +/- 0.33 | -1.07 +/- 0.36 | 0.03 +/- 0.52<br>(p = 0.9542) |
| 490-521del | -36.43 +/- 0.94 | -35.20 +/- 2.47 | 1.23 +/- 2.95<br>(p = 0.6462) | -70.86 +/- 0.43 | -70.00 +/- 0.49 | 0.86 +/- 0.71<br>(p = 0.2574) | -1.47 +/- 0.28 | -1.42 +/- 0.19 | -0.04 +/- 0.37<br>(p = 0.9206) |
| 522-555del | -32.17 +/- 1.52 | -31.25 +/- 1.10 | 0.92 +/- 2.04<br>(p = 0.6507) | -71.83 +/- 1.67 | -69.25 +/- 0.23 | 2.58 +/- 1.85<br>(p = 0.1307) | -2.42 +/- 0.49 | -2.71 +/- 0.60 | 0.29 +/- 0.83<br>(p = 0.7454) |
| 556-607del | -39.33 +/- 1.35 | -33.00 +/- 2.42 | 6.33 +/- 3.00<br>(p = 0.0691) | -74.00 +/- 0.41 | -70.29 +/- 0.94 | 3.71 +/- 1.11<br>(p = 0.0093) | -2.67 +/- 0.62 | -0.90 +/- 0.12 | -1.77 +/- 0.69<br>(p = 0.0189) |
| 608-650del | -40.14 +/- 1.19 | -35.00 +/- 1.03 | 5.14 +/- 1.70<br>(p = 0.0128) | -74.57 +/- 0.99 | -73.00 +/- 0.47 | 1.57 +/- 1.18<br>(p = 0.2355) | -1.59 +/- 0.46 | -1.18 +/- 0.35 | -0.40 +/- 0.63<br>(p = 0.5465) |
| 651-684del | -28.20 +/- 3.50 | -29.14 +/- 1.97 | -0.94 +/- 4.46<br>(p = 0.8240) | -67.00 +/- 2.48 | -68.71 +/- 0.69 | -1.71 +/- 2.87<br>(p = 0.5029) | -0.90 +/- 0.29 | -1.59 +/- 0.48 | 0.70 +/- 0.61<br>(p = 0.3286) |

Due to the low predicted disorder in the N-terminal and C-terminal regions, we suspected they might be partially or fully ordered. Checking the available Nav1.5 structures^28,29^, we noticed that portions of each region directly connected to the transmembrane domains were resolved. Consistent with this, our oocyte recordings for the deletion of the full C-terminal region resulted in current levels that were almost nonexistent, which suggested that the channel may have been too damaged either to reach the membrane or conduct current. As a result, we chose to delete only those residues that remained unresolved in the currently available structures. The partial N-terminal (430 – 457) region deletion shows a statistically significant SSI shift of −7.40 +/-2.91 and a reduction in peak current of −3.40 +/- 0.98 (**Fig. 5d-e and Table 2**). The partial C-terminal (685-698) region deletion shows a statistically significant SSI shift of −3.66 +/- 1.11 mV and change to the slow time constant of the recovery of −8.92 +/- 3.50 ms (**Fig. 5d and S1a**). Similar to 458 – 498del, the effect on the time constant did not appear to visibly alter the recovery plot (**Fig. S1e**).

**Figure 5.**
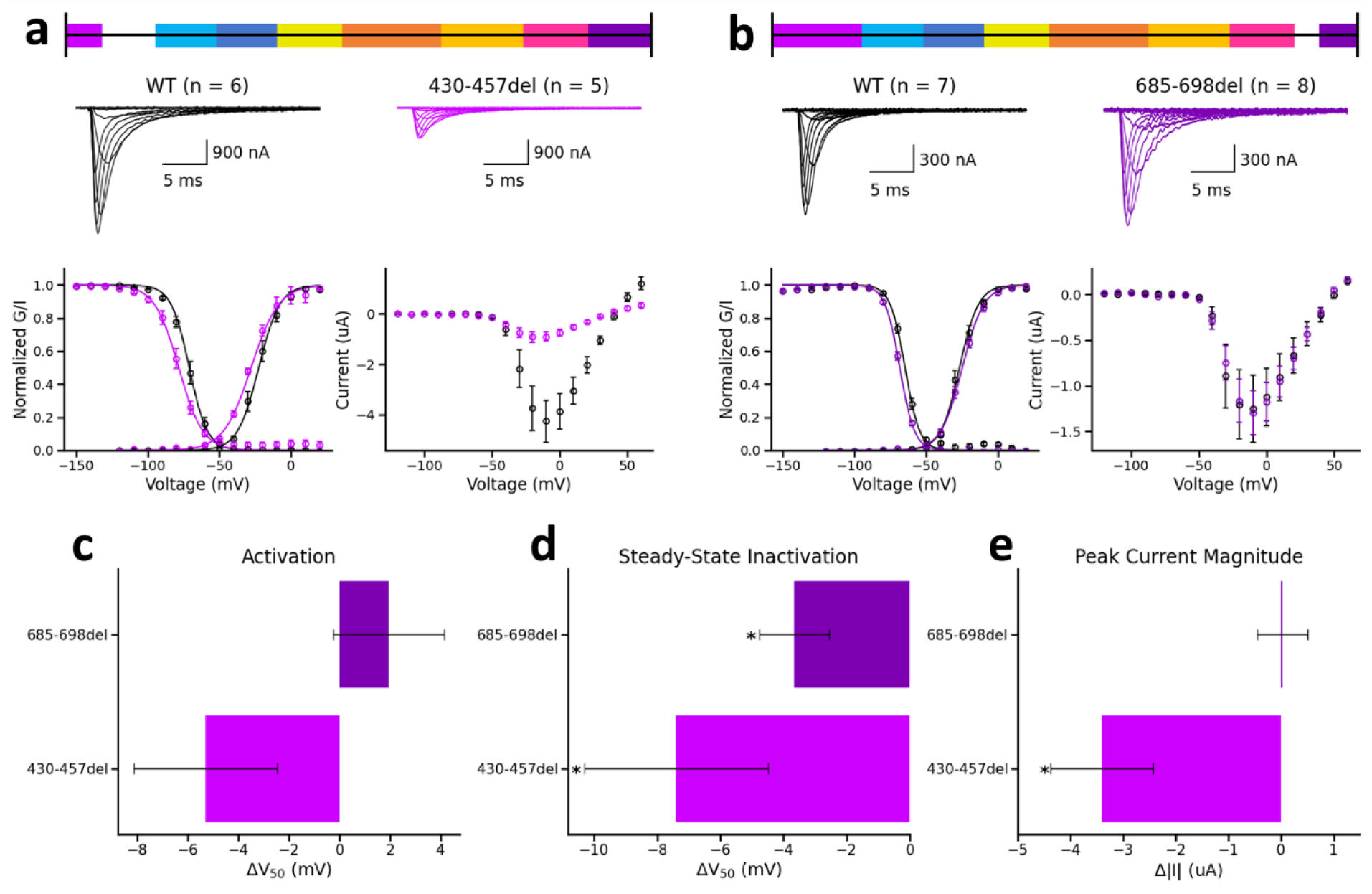
Deletions were made for the partial N-terminal (430 – 457) and partial C-terminal (685 – 698) regions. Xenopus oocyte data for deletions compared to wild-type data collected on the same days. (a – b) Xenopus oocyte data for each variant compared to wild-type data collected on the same days. For each deletion, there are representative traces for the wild-type (black) and variant (color) channels at the top of the section. Below the wild-type trace, the mean (+/- SEM) normalized conductance (G) and current (I) are plotted as functions of membrane potential and fit with single Boltzmann equations, representing channel activation and steady-state inactivation (SSI), respectively. Below the deletion trace, the mean (+/- SEM) peak current is plotted as a function of membrane potential. (c) Difference between the means (+/- SEM) of the variant and wild-type activation V_50_ from the Boltzmann fit (* indicates p-value < 0.05 and ** indicates p-value < 0.005). (d) Difference between the means (+/- SEM) of the variant and wild-type SSI V_50_ from the Boltzmann fit (* indicates p-value < 0.05). (e) Difference between the means (+/- SEM) of the variant and wild-type peak current magnitude (* indicates p-value < 0.05).

**Table 2.** Activation (GV) V_50_, Steady-State Inactivation (SSI) V_50_, and peak current magnitude (Peak, |I|) values for the I-II linker partial termini deletions.

| Construct | WT GV $V_{50}$ | Variant GV $V_{50}$ | GV $\Delta V_{50}$ (mV) | WT SSI $V_{50}$ | Variant SSI $V_{50}$ | SSI $\Delta V_{50}$ (mV) | WT Peak | Variant Peak | $\Delta \vert I\vert $ (uA) |
| --- | --- | --- | --- | --- | --- | --- | --- | --- | --- |
| 430-457del | -22.50 ± 1.97 | -27.80 ± 1.66 | -5.30 ± 2.84<br>(p = 0.1022) | -71.00 ± 1.73 | -78.40 ± 1.97 | -7.40 ± 2.91<br>(p = 0.0307) | -4.34 ± 0.87 | -0.94 ± 0.19 | -3.40 ± 0.98<br>(p = 0.0112) |
| 685-698del | -26.57 ± 1.64 | -24.62 ± 1.21 | 1.95 ± 2.19<br>(p = 0.3830) | -64.71 ± 0.83 | -68.38 ± 0.61 | -3.66 ± 1.11<br>(p = 0.0050) | -1.26 ± 0.37 | -1.29 ± 0.24 | 0.03 ± 0.48<br>(p = 0.9454) |

### Phylogenetic analyses identified several prolines as residues of interest

Conservation of specific amino acid sequences or sequence properties could suggest functional conservation, so we performed phylogenetic analyses on Nav1.5. Generally, IDRs are less conserved than their structured counterparts in terms of primary sequence conservation. However, recent studies suggest that IDRs can conserve specific features, such as residue content, net charge, isoelectric points, short linear motifs, and repeats, which can impart functional conservation.^65–68^

We ran an NCBI Protein database search for SCN5A orthologs and found sequences for more than 300 species. The majority of these species were mammalian, but there were also a number of archelosaurs, mostly birds. There were also a few reptiles, fish, and amphibians (**see Fig. S2a for clade breakdown**). We aligned these sequences with Clustal Omega and investigated the residue makeup for each region of the I-II linker.

First, we utilized pairwise distance as a measure of relative sequence conservation. Pairwise distance is the proportion of aligned residue pairs that are not identical between two sequences. The values range from zero to one, indicating identical to completely different sequences, respectively. For each species, we calculated the pairwise distance between the sequences of its aligned domains and the human wild-type sequences. Each region of the I-II linker has a different distribution of these distances, suggesting that they have different degrees of sequence conservation (**Fig. 6**). The regions predicted to be more ordered (N-terminal, Modification, and C-terminal) show higher degrees of sequence conservation (represented by lower values indicating a closer match to the human sequence) compared to the other I-II linker regions. Predictably, non-mammalian species show a higher degree of variation relative to the human sequence, suggesting a change in sequence properties across species.

**Figure 6.**
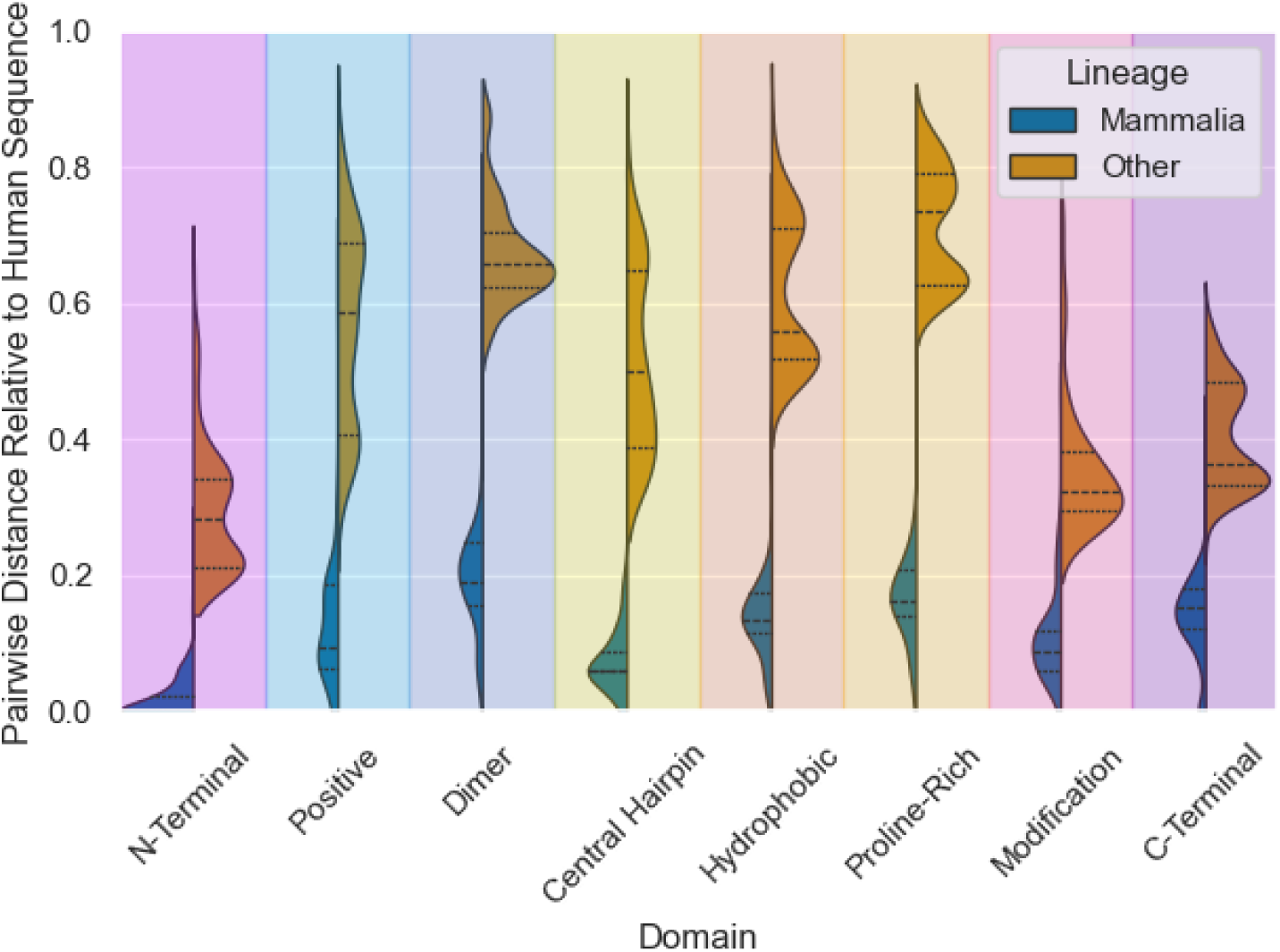
Sequence analysis reveals region- and clade-specific differences in conservation. Distribution of pairwise distances relative to the human sequence for I-II region sequences separated into mammalian and non-mammalian clades. Sequences were obtained from a search of the NCBI Protein database for eukaryotic SCN5A proteins between the lengths of 1800 and 3000 residues and aligned with Clustal Omega. For each species, the pairwise distance was calculated between the sequences of its aligned I-II regions and the human wild-type sequences. Pairwise distance is the proportion of aligned residue pairs that are not identical between two sequences. The values range from zero to one, indicating completely identical or completely different sequences, respectively.

To tease out what aspects of the sequence might be important for function within the I-II linker regions, we examined the conservation of different features (**Fig. S3**). Most notably, we found that while a relatively high proline fraction appears to be reasonably well conserved in mammals, non-mammalian sequences trend towards higher variability and lower proline fractions (**Fig. 7**). In a notable individual example, the *Xenopus* sequence is markedly proline deficient when compared to the human sequence. In our alignment, up to five proline residues that are present in humans are absent in *Xenopus* (**Fig. 7c**). Given proline’s tendency to disrupt secondary structure and promote expansion, their depletion suggests a conformational change from a more expanded conformation in mammals to a more compact conformation in non-mammals such as *Xenopus*. Thus, we wondered whether such a conformational change could confer altered channel activation.

**Figure 7.**
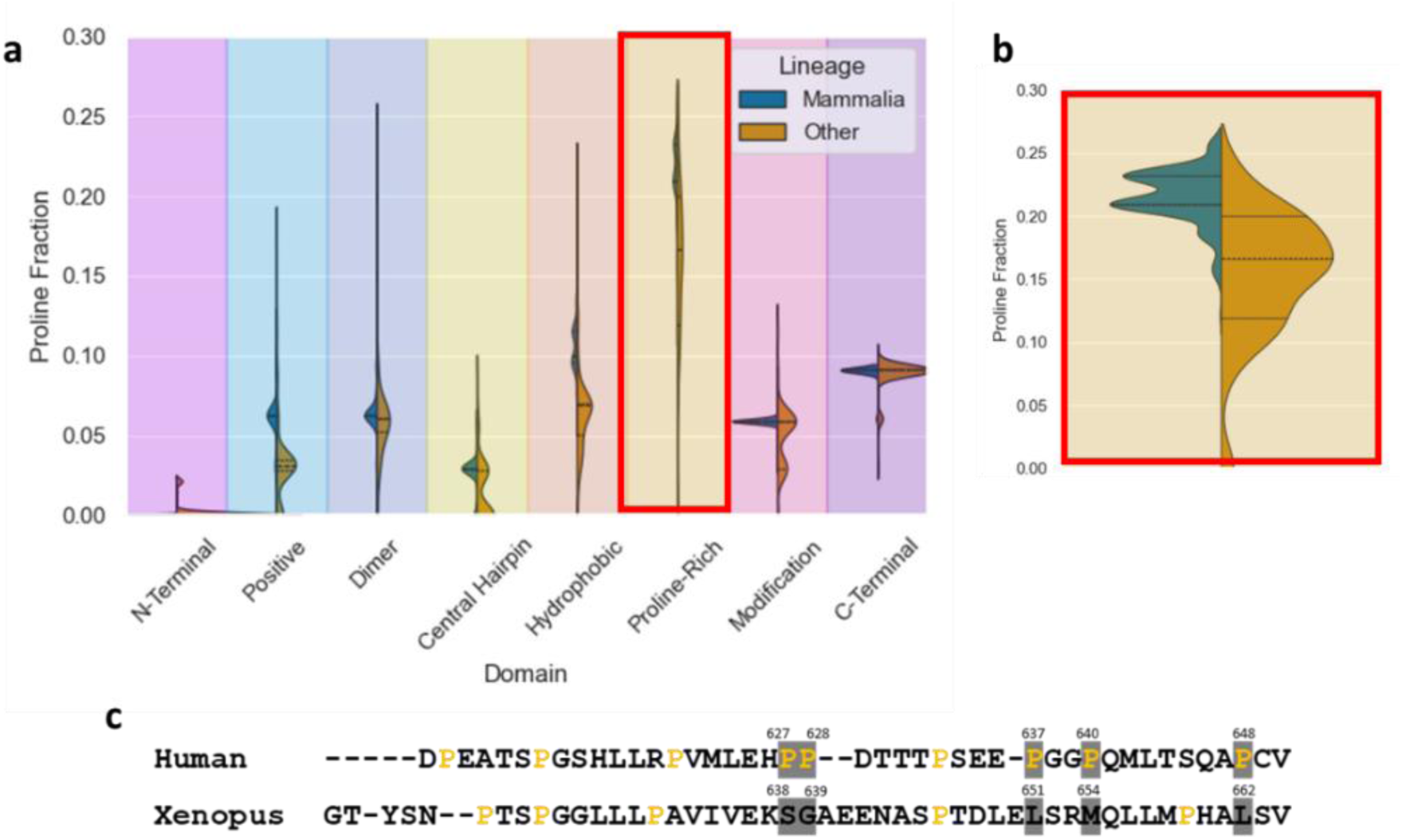
Sequence analysis reveals clade-specific differences in proline content. (a) Distribution of proline fractions for I-II region sequences separated into mammalian and non-mammalian clades. Sequences were obtained from a search of the NCBI Protein database for eukaryotic SCN5A proteins between the lengths of 1800 and 3000 residues and aligned with Clustal Omega. (b) Expanded view of proline fraction distribution for the Proline-rich region. Mammalian sequences appear to conserve high proline content in this region, whereas non-mammalian sequences are more variable. (c) Alignment of human and Xenopus Proline-rich region sequences.

### Replacing a proline with its Xenopus counterpart altered channel activation

In general, proline content is higher in the intrinsically disordered proteome than in the ordered proteome. Unlike other amino acids, a proline cannot act as a proton donor in hydrogen bonds. Their nitrogen is attached to one less hydrogen atom than it is in other residues.^61^ This property allows prolines to disrupt most standard secondary structures and increase conformational expansion, promoting disorder. A high number of prolines in a disordered protein sequence can suggest a preference for expanded conformations. The large difference in proline content between the human and Xenopus Proline-rich region could reflect significant conformational differences and the potential for functional effects. However, introducing a small difference in the proline quantity between any two sequences, such as only a single point mutation or two, is unlikely to show a noticeable difference. As a result, we started with constructs that replaced most of the prolines in the human sequence with their Xenopus counterparts to test this direction before proceeding with further experiments (Fig. 8 and Table 3).

**Figure 8.**
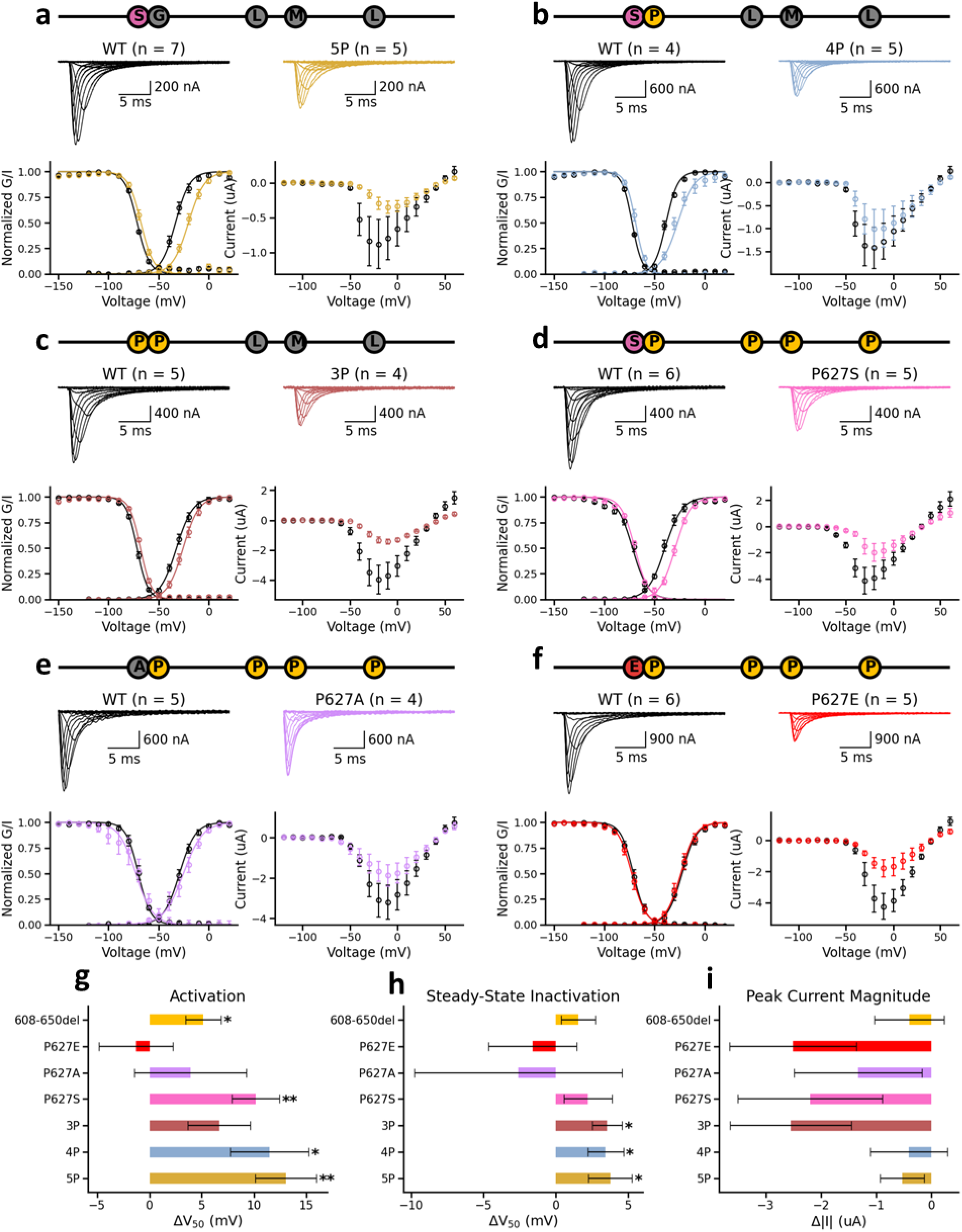
The P627S mutation depolarizes Nav1.5 activation. Variants were made and compared to the Proline-rich region (608 – 650) deletion. Xenopus oocyte data for the (a) 5P (P627S/P628G/P637L/P640M/P648L), (b) 4P (P627S/P637L/P640M/P648L), (c) 3P(P637L/P640M/P648L), (d) P627S, (e) P627A, (f) P627E variants compared to wild-type data collected on the same days. For each deletion, there are representative traces for the wild-type (black) and variant (blue) channels at the top of the section. Below the wild-type trace, the mean (+/- SEM) normalized conductance (G) and current (I) are plotted as functions of membrane potential and fit with single Boltzmann equations, representing channel activation and steady-state inactivation (SSI), respectively. Below the deletion trace, the mean (+/- SEM) peak current is plotted as a function of membrane potential. (g) Difference between the means (+/- SEM) of the variant and wild-type activation V_50_ from the Boltzmann fit (* indicates p-value < 0.05 and ** indicates p-value < 0.005). The P627S, 4P, 5P variants shows moderately sized and statistically (p- value < 0.05) significant shifts (P627S: 10.13 +/- 2.28 mV, 4P: 11.45 +/- 3.74 mV, 5P: 13.03 +/- 2.94 mV). Comparatively, deleting the entire proline-rich region results in a shift of 5.14 +/- 1.70 mV and neither the P627A nor the 3P variant exhibit a statistically significant shift. (h) Difference between the means (+/- SEM) of the variant and wild-type SSI V_50_ from the Boltzmann fit (* indicates p-value < 0.05). Like the region deletion, neither P627S nor P627S show statistically (p-value < 0.05) significant shifts. The 3P, 4P, and 5P variants all show statistically (p-value < 0.05) significant shifts. However, their small size (3P:3.55 +/- 1.02, 4P: 3.45 +/- 1.23 mV, 5P: 3.77 +/- 1.51 mV) makes functional relevance uncertain. (i) Difference between the means (+/- SEM) of the variant and wild-type peak current magnitude (* indicates p-value < 0.05). Similar to the proline-rich region deletion, none of the variants have a statistically significant effect on peak current magnitude.

**Table 3.** Activation (GV) V_50_, Steady-State Inactivation (SSI) V_50_, and peak current magnitude (Peak, |I|) values for the proline mutations.

| Construct | WT GV V<sub>50</sub> | Variant GV V<sub>50</sub> | GV Δ V<sub>50</sub> (mV) | WT SSI V<sub>50</sub> | Variant SSI V<sub>50</sub> | SSI Δ V<sub>50</sub> (mV) | WT Peak | Variant Peak | Δ \|I\| (uA) |
| --- | --- | --- | --- | --- | --- | --- | --- | --- | --- |
| 5p | -33.43 +/- 1.92 | -20.40 +/- 1.87 | 13.03 +/- 2.94<br>(p = 0.0015) | -71.57 +/- 0.34 | -67.80 +/- 1.31 | 3.77 +/- 1.51<br>(p = 0.0152) | -0.88 +/- 0.36 | -0.35 +/- 0.09 | -0.53 +/- 0.40<br>(p = 0.2869) |
| 4p | -38.25 +/- 0.74 | -26.80 +/- 3.25 | 11.45 +/- 3.74<br>(p = 0.0293) | -72.25 +/- 0.22 | -68.80 +/- 1.07 | 3.45 +/- 1.23<br>(p = 0.0408) | -1.42 +/- 0.46 | -1.01 +/- 0.42 | -0.41 +/- 0.70<br>(p = 0.5773) |
| 3p | -32.40 +/- 1.87 | -25.75 +/- 1.85 | 6.65 +/- 2.99<br>(p = 0.0636) | -71.80 +/- 0.72 | -68.25 +/- 0.54 | 3.55 +/- 1.02<br>(p = 0.0124) | -3.99 +/- 0.97 | -1.44 +/- 0.16 | -2.55 +/- 1.10<br>(p = 0.0790) |
| p627s | -39.33 +/- 1.74 | -29.20 +/- 1.11 | 10.13 +/- 2.28<br>(p = 0.0022) | -71.83 +/- 1.44 | -69.60 +/- 0.46 | 2.23 +/- 1.66<br>(p = 0.2475) | -4.20 +/- 0.99 | -2.00 +/- 0.66 | -2.20 +/- 1.31<br>(p = 0.1434) |
| p627a | -29.40 +/- 2.01 | -25.50 +/- 4.22 | 3.90 +/- 5.37<br>(p = 0.4595) | -70.40 +/- 1.82 | -73.00 +/- 5.96 | -2.60 +/- 7.18<br>(p = 0.6998) | -3.26 +/- 0.87 | -1.93 +/- 0.54 | -1.33 +/- 1.16<br>(p = 0.3171) |
| p627e | -22.50 +/- 1.97 | -23.80 +/- 2.52 | -1.30 +/- 3.55<br>(p = 0.7178) | -71.00 +/- 1.73 | -72.60 +/- 2.13 | -1.60 +/- 3.04<br>(p = 0.6070) | -4.34 +/- 0.87 | -1.83 +/- 0.57 | -2.51 +/- 1.15<br>(p = 0.0663) |

First, we created two Nav1.5 constructs where we replaced either four (4P, P627S/P637L/P640M/P648L) or five (5P, P627S/P628G/P637L/P640M/P648L) of the prolines with their Xenopus counterparts. Proline mutation choices were made based on the alignment between the human and *Xenopus* sequence. We observed a moderately-sized, statistically significant activation shift for both variants (4P: 11.45 +/- 3.74 mV, 5P: 13.03 +/- 2.94 mV) that was approximately twice the size of the shift seen when the entire proline-rich region was removed (**Fig. 8g**). Both variants also showed SSI shifts (**Fig. 8h**) that, while statistically significant, were quite small (4P: 3.45 +/- 1.23 mV, 5P: 3.77 +/- 1.51 mV). Additionally, neither variant had a statistically significant effect on peak current magnitude (**Fig. 8i**), though the 5P plot appears to show one (**Fig. 8d**); and neither altered the recovery time constants or the late current (**Fig. S1a-c**).

Next, we created a construct that replaced only three of the prolines (3P, P637L/P640M/P648L) and did not contain P627S, since the introduction of a serine residue could have effects that are unrelated to proline content. This variant retained the small SSI shift (3.55 +/- 1.02) (**Fig. 8h**), the seemingly visible but not statistically significant change in the peak current magnitude (**Fig. 8d and** **8i**), and the lack of effects on the slow recovery time constant and the late current seen in the 5P variant. However, this variant did not retain the activation shift nor the lack of statistically significant effect on the fast recovery time constant (−0.39 +/- 0.12 ms). Unlike the two deletions with an effect on the slow time constant, this alteration of the recovery was visible in the plot (**Fig. S1b and S1f**).

To determine whether altered channel activation was due to a reduction in proline content or to the effects of a point mutation, we focused on mutations to P627. In the 4P and 5P constructs, P627 was mutated to a serine which could lead to a new phosphorylation site within the proline-rich region. Thus, we created three constructs where we replaced only the proline at position 627. The first, P627S, corresponded to the aligned Xenopus residue, the second, P627A, served as a reference point for the potential serine phosphorylation, and the third, P627E, as a phosphomimetic mutation. Unlike the 4P and 5P constructs, none of these variants affected the SSI (**Fig. 8h**). Further, neither the recovery time constants nor the late current were affected by the P627 variants (**Fig. S1**). Only P627S resulted in a statistically significant (p < 0.05) effect: a 10.13 +/- 2.28 mV depolarizing activation V_50_ shift (**Fig. 8a**), similar in size to those seen for both the 4P and the 5P variants. Neither P627A nor P627E had a statistically significant effect (**Fig. 8b**).

## Discussion

The DI-DII linker of Nav1.5 has so far been associated with many different channel functions, but its definitive role is not well understood. An in-depth look at its sequence revealed a series of regions with distinct properties. We theorized that these regions could have distinct roles in channel function and investigated by observing channel function when each region was removed. Despite removing such large portions of the channel, some of which are well-conserved across species and channel isoforms, we observed negligible effect on channel gating, even though peak current magnitude was significantly altered, suggesting that the primary role of the DI-DII linker could be theregulation of channel expression. Unlike with mammalian cell transfection, the RNA injection technique used to express the channels in oocytes allows us a relatively tight control over the amount of nucleic material entering the cell. As a result, changes in the peak current magnitude can suggest that alterations to the cell surface expression have occurred. However, it is important to note that many of the physiological processes in oocytes differ greatly from those in mammalian cells.^69^ As such, it may be difficult to extrapolate changes in channel expression and trafficking in oocytes to functional relevance in the native cardiomyocytes. Further, oocytes lack many of the regulatory, co-expressing, and interacting proteins found in the channel’s native cardiomyocyte environment.^69^ While this does allow the channel and components to be studied in isolation, it also limits our ability to evaluate the physiological relevance of our results. The lack of effect the deletions had on channel gating may actually suggest the absence of important interacting partners.

Both the 430 – 457del (partial N-terminal deletion) and 556 – 607del (Hydrophobic region deletion) resulted in conspicuous decreases in peak current magnitude, which could imply a decrease in cell surface expression and may suggest a role for the linker in trafficking and membrane retention rather than gating. Intriguingly, 430 – 457del includes several interaction sites associated with channel expression. The most notable is NEDD4L ubiquitination, which can decrease peak current density and cell surface expression.^47^ Four ubiquitination sites were identified in the DI-DII linker, three of which (K430, K442, and K443) are covered by this deletion. Furthermore, this deletion overlaps part of a dynamitin interaction (417 – 444), which was also shown to decrease current amplitude and cell surface density.^70^ Additionally, the L567Q variant, which has been associated with a reduction in current density, resides within 556 – 607del.^71^ Unfortunately, it is unclear how this information fits with the decreases in current we observed for the deletions, in particular for 430 – 457del. Since ubiquitination is known to decrease current density and expression, we might expect the removal of 430 – 457 to increase the current by removing K430, K442, and K443 and reducing ubiquitination. However, it is also possible that the channel does not undergo any significant ubiquitination in our experiments due to the use of the oocyte expression system. Endogenous NEDD4 has been found in Xenopus oocytes previously but its expression levels appear to be relatively low. ^72,73^ It may not be at high enough levels to play a significant role in our experiments. In this case, the effect of the deletion of 430 – 457 could be dominated by another mechanism. For example, if the three lysines (K430, K442, and K443) increase expression when they are not ubiquitinated, their removal could have a similar effect as their ubiquitination.

Interestingly, the skeletal muscle isoform, Nav1.4, functions normally without many of the regions we deleted. It is approximately 180 residues shorter than Nav1.5 and 173 of those residues are in the DI-DII linker. Of the eight DI-DII regions, Nav1.4 only retains three: the N-terminal, the Modification, and the C-terminal domains. The 2004 Bennet study suggested that the disordered linkers may play a role in activation.^37^ In that study, chimeras were made of Nav1.5 and Nav1.4 where the I-II and II-III linkers were switched between the isoforms, both individually and together, and recorded in CHO cells. The substitution of Nav1.4 I-II into Nav1.5 is roughly equivalent to the combined deletion of the Positive, Dimer, Hairpin, Hydrophobic, and Proline-rich regions, yet it only has a relatively small effect on activation (5.8 mV hyperpolarizing shift), on the same order of magnitude as the depolarizing shift (5.14 mV) we observed when we removed the Proline-rich region. The relatively small impact on channel function observed for this substitution is consistent with the lack of gating effect we observed for the Positive, Dimer, Hairpin, Hydrophobic, and Proline-rich region deletions, suggesting that the primary functional differences between the two isoforms may be complex, possibly involving protein or environmental interactions.

Interestingly, the only constructs that altered channel gating significantly (P627S, P627S/P637L/P640M/P648L, P627S/P628G/P637L/P640M/P648L) all included the same mutation: P627S. All three of these constructs had similar effects on channel activation, suggesting that the introduction of the serine was responsible for the changes in channel activation observed in the three original constructs. Combined with the lack of effect observed for the P627A and P627E mutations, these results suggest that the activation shift may depend on a serine-specific property. The most obvious property to consider would be serine’s ability to undergo phosphorylation, though this would require further experiments to determine. Despite the relative lack of potential interaction partners in oocytes, the cells are not empty and do have their own endogenous proteins, including some kinases, which could be responsible for this gating effect. Adding a serine at position 627 creates a sequence that aligns with the known binding motifs of both MAPK (LEH<u>S</u>PDT) and GSK3 (LEH<u>S</u>PDTT), which would phosphorylate the serine at 627. A CK1 motif (SPD<u>T</u>TTP) was also created that would phosphorylate threonine at 630 instead of the mutant serine at 627.

The Modification region (651 – 684) contains a cluster of phosphorylation sites implicated in the regulation of channel activation and current density.^49^ Replacing either S664 or S667 with either alanine or glutamate depolarized channel activation by approximately 6 mV. Since both the phosphosilent and the phosphomimetic mutations for S664 and S667 had the same effect on channel gating, phosphorylation at these sites may be the default state and the activation shift may be due to the disruption of that interaction. P627S may present an interesting comparison. Neither P627A nor P627E have any effect on channel gating, yet P627S causes a noticeable depolarizing shift. If the wild type residue at position 627 was a serine instead of a proline, the channel might show a response to phosphosilent and phosphomimetic mutations in the same vein as those seen for S664 and S667.

The impressive resistance of channel gating to the removal of large regions combined with the presence of many known interaction sites could imply that the I-II linker’s fundamental role is as an interface for interactions with other proteins. In this case, these proteins may need to be present for a deletion to impact gating. It is likely that, without the interacting proteins present, removing the regions has little effect on channel function. Consequentially, variants may have a larger impact if they disrupt these interactions. Determining whether they do so may be helpful in evaluating the likely pathogenicity of variants. For example, the Hydrophobic region (556 – 607) contains a residue, S571, that has been reported to increase the late sodium current when phosphorylated by CaMKII.^50^ The mutations, A572D and Q573E, have been shown to alter the CaMKII-dependent regulation of Nav1.5 and partially mimic the effect of S571 phosphorylation on channel function, with potentially pathogenic consequences.^74^

The Proline-rich region (608 – 650) has a possible SH3-binding motif (XXXPXXP) that includes the P637 and P640 residues. This binding motif could act as an anchoring site for the SH3 domains of proteins that interact with other areas of the channel, in which case the effect of region deletion would likely only be apparent in a more native-like environment containing those proteins. Under native conditions, the disruption of this anchor site by the variants of unknown significance, P637L ^35^ and P640A ^34^,could greatly impact channel function. It may be difficult to truly evaluate the pathogenicity of these variants without the correct endogenous proteins, however.

The Positive (458 – 489) and Dimer (490 – 521) regions are primarily associated with the dimerization interaction from Clatot et al. 2017.^27^ This interaction may itself require clustering/localization within the cell as well as certain partner proteins, such as 14-3-3. There are many variants of unknown significance in these regions that could act pathogenically through the disruption of either the alpha-alpha or other protein interactions. Additionally, the Dimer region contains an arginine methylation site (R513)^52^ and a PKA phosphorylation site (S516).^51^ Interestingly, the R513 methylation and S516 phosphorylation interactions appear to block each other. ^51^The methylation, but not the phosphorylation is abolished by the disease-associated mutation, G514C.^51^

Finally, the Hairpin region (522 – 555) contains an arginine methylation site (R526)^52^ and two PKA phosphorylation sites (S525 and S528). The PKA sites are associated with a peak current increase through the mediation of nearby ER-retention motifs.^44,48^ Both the R526H and S528A mutations disable PKA phosphorylation and reduce channel expression.^44^ Further, this region contains a MOG1 interaction domain (F530 – R535), which has been connected to trafficking and an increase in current amplitude. In Xiong et al. 2022, the F530A, F532A, R533A, and R534A mutations all reduced MOG1 interaction and eliminated the increased current, as did the disease-associated mutation F532C.^75^ Though another disease-associated variant, R535Q, reduced current amplitude in a MOG1-independent manner and a further one, F530V, had no apparent effect, this still provides support for the idea that mutations that disrupt interaction sites are more interesting candidates for possible pathologies.

Future studies may focus on identifying and characterizing I-II linker interactions. We have found that the deletion of large regions from the linker has little effect on channel gating. Comparatively, small changes in the sequence at key positions seem to have a higher impact, particularly if they create or disrupt interaction sites. *Xenopus* oocytes provide a useful system if we want to study the effect of individual interaction partners. Studies in native myocytes, may reveal interacting proteins that are required for the DI-DII linker to further regulate channel function.

