## Supplemental Data for "Investigating the Functional Role of the DI-DII Linker in Nav1.5 Channel Function"

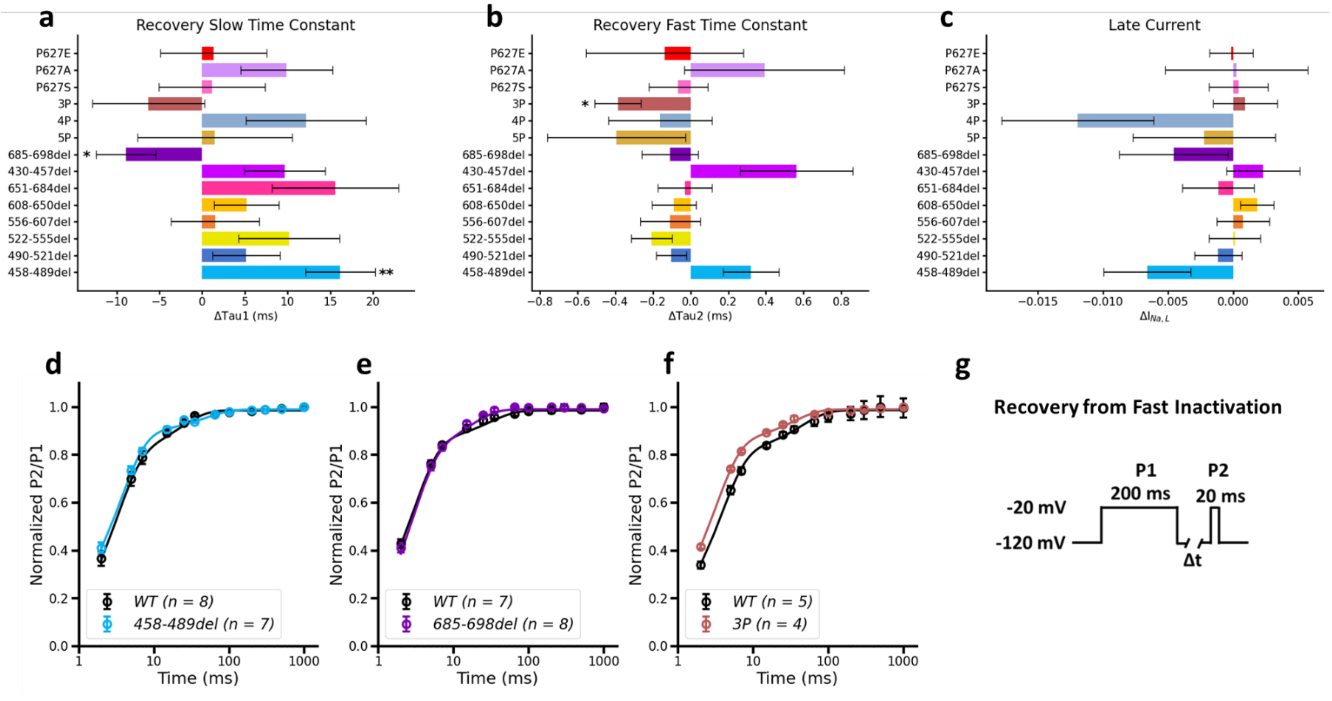

**Figure S1. Neither deletions nor proline replacements significantly alter either the recovery from inactivation or the late current.** Includes data for the Positive (458 – 489), Dimer (490 – 521), Hairpin (522 – 555), Hydrophobic (556 – 607), Proline-rich (608 – 650), Modification (651 – 684), partial N-terminal (430 – 457), and the partial C-terminal (685 – 698) region deletions as well as the 3P (P637L/P640M/P648L), 4P (P627S/P637L/P640M/P648L), 5P (P627S/P628G/P637L/P640M/P648L), P627S, P627A, and P627E variants. (a) Difference between the means (+/- SEM) of the variant/deletion and wild-type slow time constant of the recovery from inactivation (* indicates p-value < 0.05 and ** indicates p-value < 0.005). The Positive and partial C-terminal region deletions show the only statistically (p-value < 0.05) significant shifts. However, the small size (Positive: 16.16 +/- 4.07 ms, partial C-terminal: -8.92 +/- 3.50 ms) of these shifts makes their functional relevance uncertain. (b) Difference between the means (+/- SEM) of the variant/deletion and wild-type fast time constant of the recovery from inactivation. Only the 3P variant shows a statistically significant change (-0.39 +/- 0.12 ms). (c) Difference between the means (+/- SEM) of the variant/deletion and wild-type late current. (d – f) Recovery curves for the three constructs that showed a statistically significant effect on either the slow time constant (458 – 489del and 685 – 698del) or the fast time constant (3P). (g) Stimulation protocol used to obtain the recovery from fast inactivation.

| Construct | WT Tau1 | Variant Tau1 | Δ Tau1 (ms) | WT Tau2 | Variant Tau2 | Δ Tau2 (ms) | WT $I_{Na,L}$ | Variant $I_{Na,L}$ | Δ $I_{Na,L}$ |
|---|---|---|---|---|---|---|---|---|---|
| 458-489del | 14.85 +/- 0.94 | 31.01 +/- 3.66 | 16.16 +/- 4.07 (p = 0.0010) | 2.02 +/- 0.12 | 2.34 +/- 0.08 | 0.32 +/- 0.15 (p = 0.0569) | 0.0097 +/- 0.0052 | -0.0060 +/- 0.0066 | -0.0066 +/- 0.0033 (p = 0.0654) |
| 490-521del | 24.77 +/- 1.82 | 29.93 +/- 3.09 | 5.16 +/- 3.97 (p = 0.1939) | 2.21 +/- 0.04 | 2.11 +/- 0.06 | -0.10 +/- 0.08 (p = 0.2160) | 0.0007 +/- 0.0032 | 0.0039 +/- 0.0025 | -0.0012 +/- 0.0018 (p = 0.5507) |
| 522-555del | 28.50 +/- 4.04 | 38.68 +/- 3.63 | 10.18 +/- 5.89 (p = 0.1099) | 2.37 +/- 0.09 | 2.17 +/- 0.04 | -0.21 +/- 0.11 (p = 0.0576) | 0.0011 +/- 0.0042 | 0.0044 +/- 0.0017 | 0.0001 +/- 0.0020 (p = 0.9488) |
| 556-607del | 22.42 +/- 1.72 | 23.96 +/- 4.45 | 1.54 +/- 5.17 (p = 0.7843) | 2.42 +/- 0.04 | 2.31 +/- 0.14 | -0.11 +/- 0.16 (p = 0.5442) | 0.0066 +/- 0.0015 | 0.0080 +/- 0.0047 | 0.0008 +/- 0.0020 (p = 0.7320) |
| 608-650del | 22.24 +/- 1.82 | 27.42 +/- 2.96 | 5.18 +/- 3.79 (p = 0.1855) | 2.23 +/- 0.08 | 2.14 +/- 0.08 | -0.09 +/- 0.12 (p = 0.4710) | 0.0022 +/- 0.0017 | 0.0109 +/- 0.0024 | 0.0018 +/- 0.0013 (p = 0.1663) |
| 651-684del | 15.85 +/- 4.04 | 31.45 +/- 5.43 | 15.60 +/- 7.40 (p = 0.0789) | 1.98 +/- 0.09 | 1.95 +/- 0.10 | -0.03 +/- 0.14 (p = 0.8468) | 0.0071 +/- 0.0053 | 0.0068 +/- 0.0018 | -0.0011 +/- 0.0027 (p = 0.6374) |
| 430-457del | 21.62 +/- 3.90 | 31.30 +/- 1.80 | 9.68 +/- 4.73 (p = 0.0883) | 2.62 +/- 0.21 | 3.18 +/- 0.17 | 0.56 +/- 0.30 (p = 0.1000) | 0.0099 +/- 0.0019 | 0.0118 +/- 0.0054 | 0.0023 +/- 0.0028 (p = 0.4029) |
| 685-698del | 19.76 +/- 2.97 | 10.84 +/- 1.32 | -8.92 +/- 3.50 (p = 0.0194) | 1.90 +/- 0.05 | 1.79 +/- 0.13 | -0.11 +/- 0.15 (p = 0.5041) | 0.0071 +/- 0.0048 | -0.0240 +/- 0.0098 | -0.0046 +/- 0.0042 (p = 0.3140) |
| 5p | 30.12 +/- 5.37 | 31.60 +/- 6.21 | 1.48 +/- 9.05 (p = 0.8730) | 2.59 +/- 0.27 | 2.19 +/- 0.20 | -0.39 +/- 0.37 (p = 0.3426) | 0.0044 +/- 0.0079 | 0.0224 +/- 0.0089 | -0.0022 +/- 0.0055 (p = 0.6858) |
| 4p | 20.56 +/- 3.21 | 32.71 +/- 5.33 | 12.15 +/- 7.02 (p = 0.1493) | 2.17 +/- 0.07 | 2.01 +/- 0.23 | -0.16 +/- 0.28 (p = 0.6153) | 0.0065 +/- 0.0019 | -0.0121 +/- 0.0115 | -0.0120 +/- 0.0058 (p = 0.1112) |
| 3p | 28.63 +/- 4.98 | 22.36 +/- 2.96 | -6.27 +/- 6.54 (p = 0.3996) | 2.40 +/- 0.06 | 2.02 +/- 0.09 | -0.39 +/- 0.12 (p = 0.0147) | 0.0176 +/- 0.0027 | 0.0115 +/- 0.0036 | 0.0009 +/- 0.0025 (p = 0.7081) |
| p627s | 29.17 +/- 3.73 | 30.35 +/- 4.19 | 1.18 +/- 6.22 (p = 0.8536) | 2.40 +/- 0.12 | 2.34 +/- 0.08 | -0.07 +/- 0.16 (p = 0.6972) | 0.0045 +/- 0.0037 | 0.0013 +/- 0.0031 | 0.0004 +/- 0.0023 (p = 0.8630) |
| p627a | 19.77 +/- 1.33 | 29.66 +/- 4.49 | 9.89 +/- 5.39 (p = 0.0808) | 2.62 +/- 0.23 | 3.01 +/- 0.29 | 0.39 +/- 0.42 (p = 0.3756) | 0.0027 +/- 0.0033 | 0.0084 +/- 0.0090 | 0.0003 +/- 0.0055 (p = 0.9602) |
| p627e | 21.62 +/- 3.90 | 22.97 +/- 4.07 | 1.35 +/- 6.24 (p = 0.8342) | 2.62 +/- 0.21 | 2.48 +/- 0.31 | -0.14 +/- 0.42 (p = 0.7423) | 0.0099 +/- 0.0019 | 0.0091 +/- 0.0029 | -0.0002 +/- 0.0017 (p = 0.9230) |

**Table S1. Recovery Slow Time Constant (Tau1), Recovery Fast Time Constant (Tau2), and Late Current ($I_{Na,L}$) values for the deletion and proline mutations**

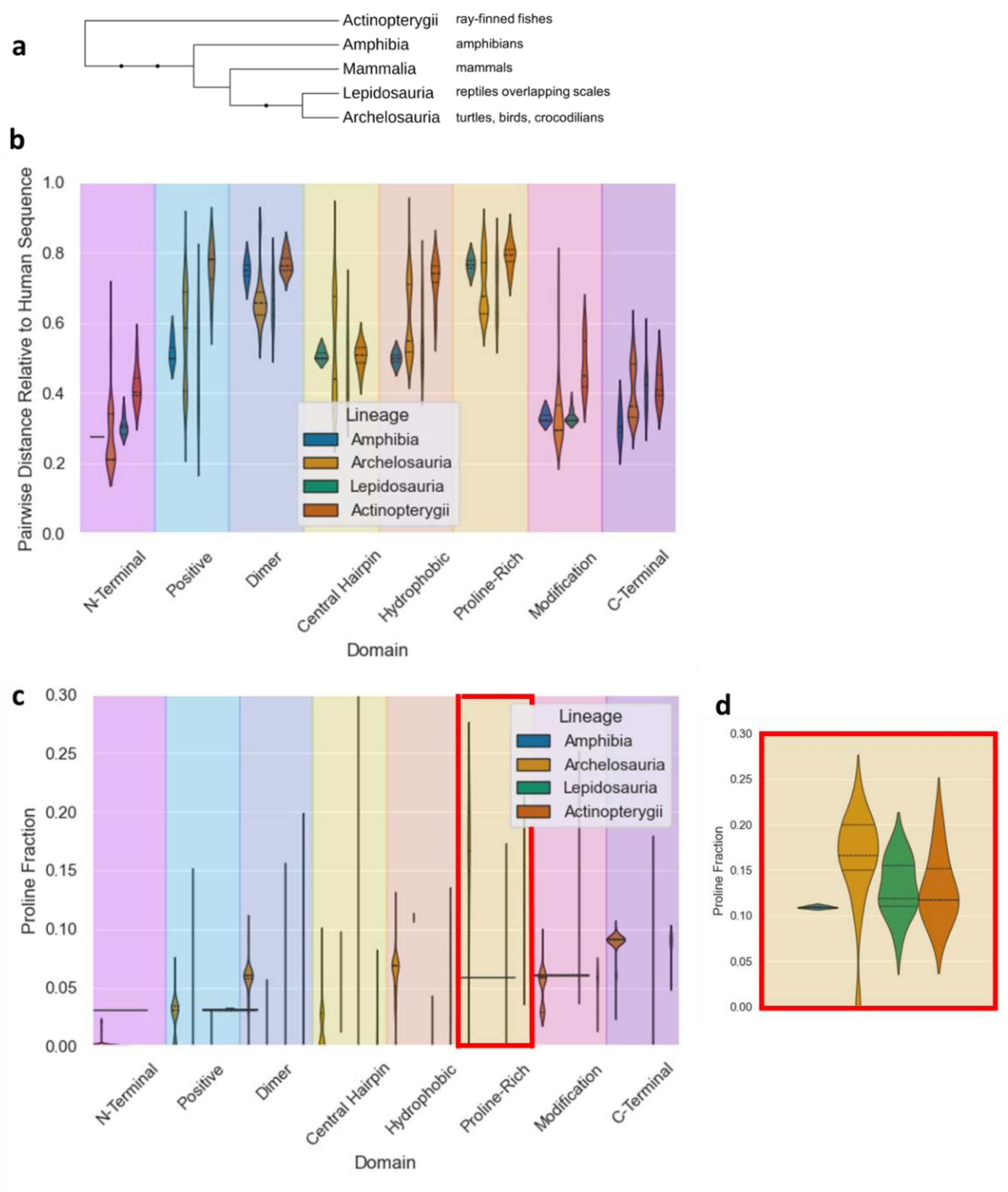


**Figure S2. Clade breakdown of proline-content for non-mammalian sequences.** (a) Clades of the sequences obtained by NCBI Protein database search for eukaryotic SCN5A proteins between the lengths of 1800 and 3000 residues. (b) Distributions of pairwise distances relative to the human sequence for non-mammalian clades. (c) Distributions of proline fractions for non-mammalian clades. (d) Expanded view of proline fraction distributions for the Proline-rich region.

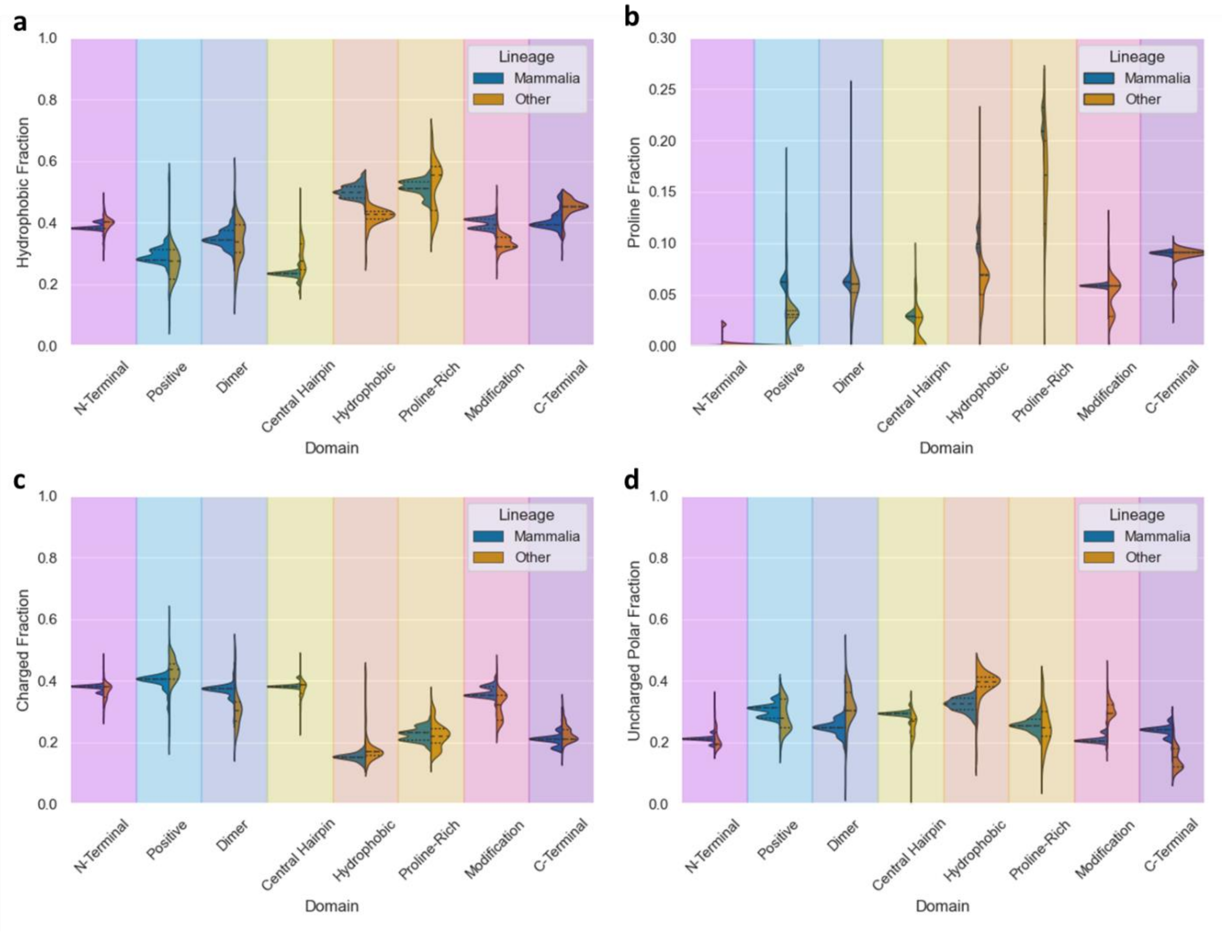


**Figure S3. Region- and clade-specific differences in I-II residue content.** Distributions of (a) hydrophobic, (b) proline, (c) charged, and (d) uncharged polar residue fractions for I-II region sequences separated into mammalian and non-mammalian clades.

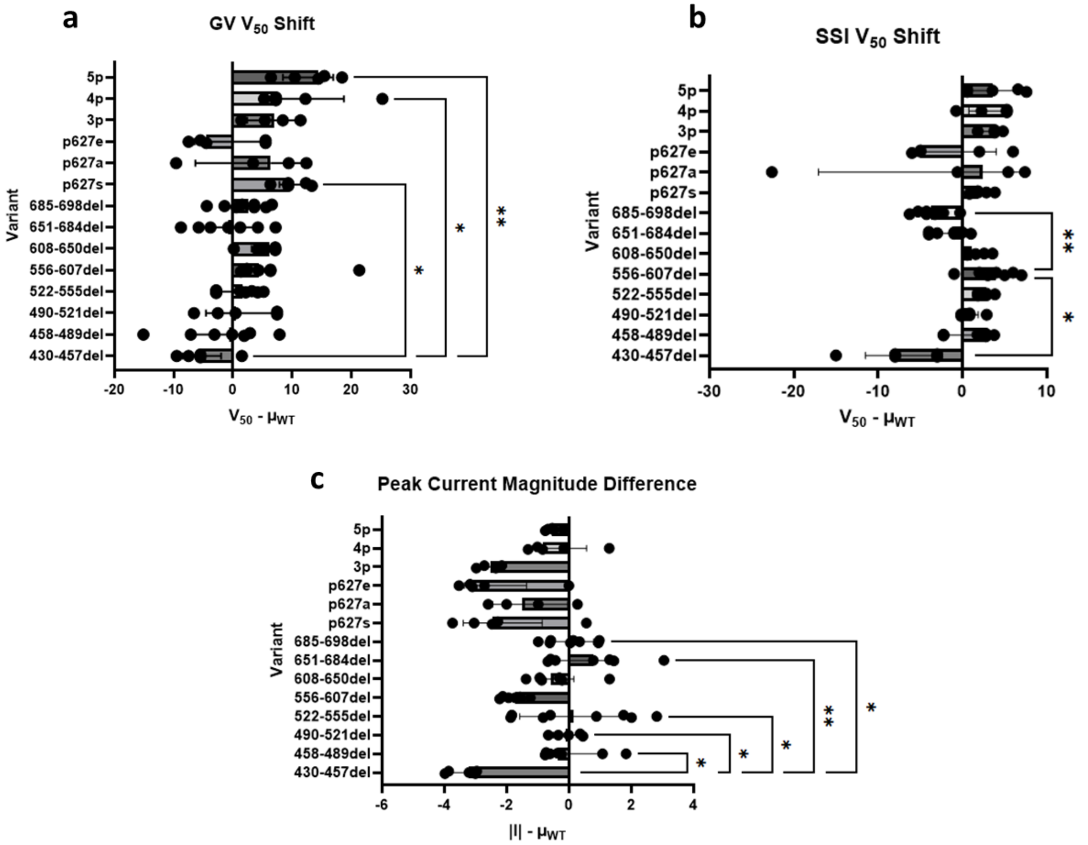


**Figure S4. ANOVA analysis of the activation, inactivation, and peak current changes.** For each variant, the average V50 or magnitude value for WT cells from the same oocyte batch recorded on the same day was subtracted from each variant V50 or magnitude value. This created distributions of differences of (a) the activation (GV), (b) the steady-state inactivation (SSI), and (c) the peak current magnitude for each variant. Then, we then ran a Kruskal-Wallis test and a Dunn's multiple comparison test, notating which comparisons were statistically significant.
